# Adaptation of fluorescence rescue reporters of mistranslation for use in *Saccharomyces cerevisiae*

**DOI:** 10.64898/2026.10.20.760511

**Authors:** Brendan S. Charles, Amanda J. Moehring

**Affiliations:** Western University Department of Biology, Western University, London, Ontario, Canada, N6A 5B7

**Keywords:** Mistranslation, tRNA, misfolded protein, proteostasis, *Saccharomyces cerevisiae*, *Drosophila melanogaster*

## Abstract

Specific mutations to tRNA genes can cause mistranslation - the incorporation of amino acids mismatched to the mRNA code during translation. The effects of mistranslation are hugely varied, depending on the form of mistranslation and various contextual factors. A key aspect of mutant tRNA mistranslation investigation is the reliable detection of mistranslation events. While mistranslation confirmation and quantification are commonly accomplished via mass spectrometry, others have devised a means of detecting mistranslation via fluorescent signal.

Here, we successfully adapt this method for use in *Saccharomyces cerevisiae*, a model in which a great deal of mistranslation research is currently conducted. We demonstrate that mistranslation fluorescence rescue is detectable using both microscopy as well as fluorescent plate reader measurement, and that fluorescence intensity varies with the form of mistranslation assessed.

Interestingly, the quantity of mistranslation measured by fluorescence rescue differs from measurements by standard mass spectrometry methods, potentially due to capturing a different timepoint of proteins that contain mistranslation errors. As such, this tool has high utility in detecting mutant tRNA mistranslation presence in yeast models, while its quantification of mistranslation frequency requires further investigation.

## Introduction

Mistranslation via mutant tRNAs can have diverse biological impacts dependent on the form of and frequency of mistranslation (Charles et al., 2026, Cozma et al., 2023; Davey-Young et al., 2024; Fan et al., 2015; Schwartz & Pan, 2016; Zimmerman et al., 2018). In order to make claims regarding the biological outcomes of mistranslation, one must be able to reliably detect and quantify mistranslation events. Most commonly, this is accomplished by the use of mass spectrometry to measure the prevalence of mutant peptides relative to their wildtype counterparts (Cozma et al., 2023; Davey-Young et al., 2024; Isaacson et al., 2024). However, mass spectrometry can be a costly and time intensive process. Additionally, more nuanced assessments of mistranslation, such as tissue or temporally specific mistranslation frequencies, complicate sample preparation for use with mass spectrometry. For these reasons, various approaches to reporting mistranslation via fluorescence have been established in multiple model organisms (Gomes et al., 2016; Lant et al., 2018; Rozik et al., 2022; Sagi et al., 2016). Despite a great deal of mistranslation research having been conducted in *Saccharomyces cerevisiae*, none of the established fluorescent reporters of mistranslation have been established in this model.

Fluorescence reporters of mistranslation function by mistranslation-dependent induction of fluorescent signal. This is accomplished by one of two mechanisms. The method chosen for adaptation here is reliant upon loss of function mutations to fluorescent proteins, which are rescued by the corresponding form of mistranslation (Figure 1)(Gomes et al., 2016; Lant et al., 2018; Rozik et al., 2022). For instance, an mCherry protein bearing an S151V mutation is rendered non-functional and thus produces no red fluorescence. When expressed alongside a mutant tRNA^Ser^ that mistranslates Ser at Val codons, the S151V mutation in mCherry is restored to a Ser residue, thus restoring function and producing red fluorescence. The second mechanisms for fluorescent reporting of mistranslation utilizes a stop codon preceding the fluorescent protein coding sequence (Sagi et al., 2016). Without the presence of a mutant mistranslating tRNA, the fluorescent protein is never translated, but in the presence of a mutant tRNA designed to cause stop codon readthrough, the fluorescent protein is expressed. With either such mechanisms, the mistranslation-dependent fluorescent signal is typically measured relative to a second fluorescent protein which serves as an expression control (Gomes et al., 2016; Sagi et al., 2016).

**Figure 1.**
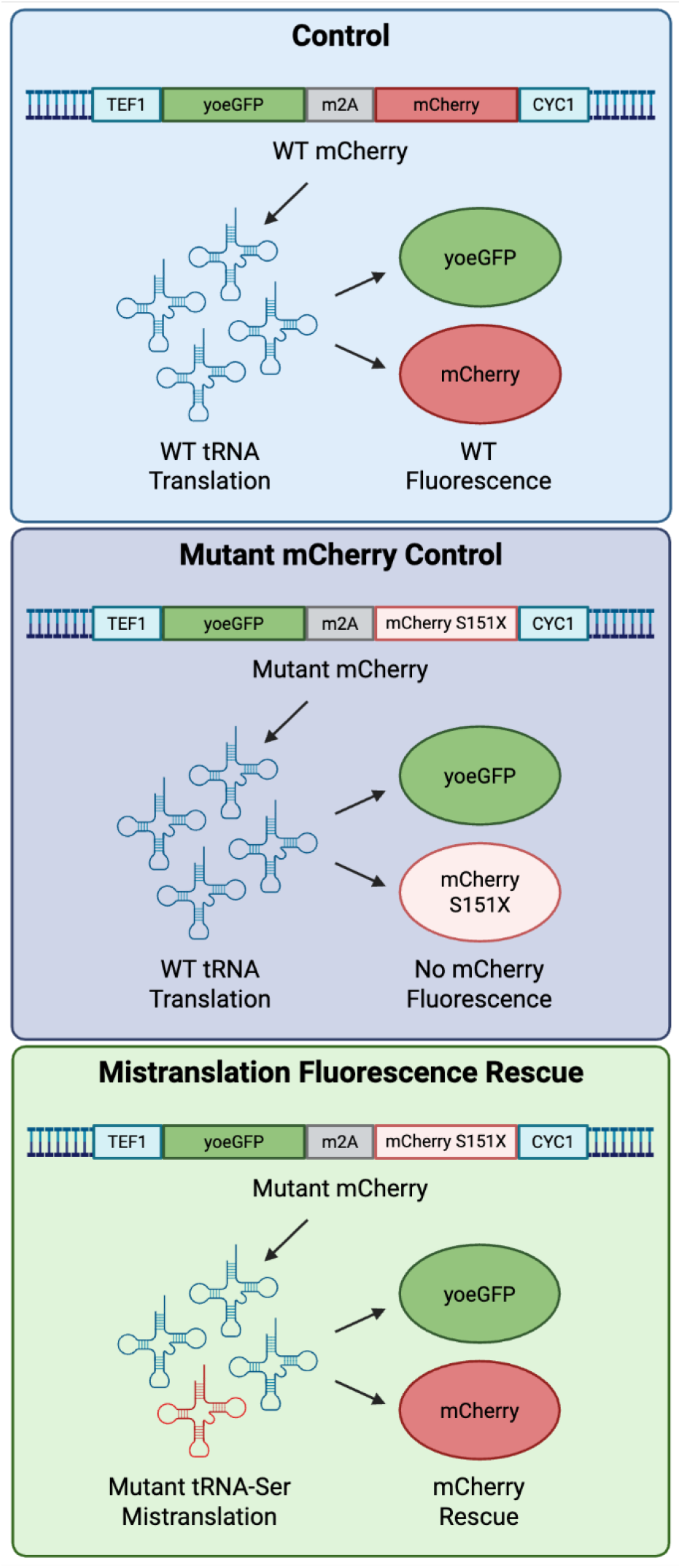
Schematic of mechanism for mistranslation fluorescence rescue. Top: A single genetic construct produces a transcript coding for both yoeGFP and mCherry; the proteins are produced separately due to the m2A separating sequence. Center: A mutation at the S151X location in mCherry prevents identification of this locus by a wildtype serine tRNA and causes loss of function of the mCherry protein; only the control fluorophore yoeGFP is functional. Bottom: Co-expression of mCherry bearing an S151X loss of function mutation alongside mutant tRNA^Ser^ designed to mistranslate the mutant codon at S151X back to a serine results in rescue of fluorescent signal.

Here, we combine mutant mistranslating tRNAs with mutations to yeast-optimized fluorescent proteins for detection of serine mistranslation in *Saccharomyces cerevisiae*. Mutation of the S151 locus of mCherry is observed to ablate fluorescence signal in the absence of mistranslation, while the co-expression of S151 mutants alongside corresponding mutant mistranslating tRNA^Ser^ restores mild red fluorescence for some forms of mistranslation (Rozik et al., 2022). Interestingly, the quantification of red fluorescence from mistranslation differs substantially from standard mass spectrometry methods of detecting and quantifying mistranslation from the same mistranslating tRNA variants (Charles & Moehring, 2026).

## Methods

### DNA cloning

The pDRF1-GW yoeGFP-T2A-mCherry plasmid containing a yeast codon optimized eGFP linked to mCherry with the self-digesting M2A peptide sequence was ordered from Addgene (#118446). The dual fluorescent protein sequence was PCR cloned 5’ SpeI and 3’XhoI into a p413 TEF yeast expression plasmid with a histidine selectable marker such that the fluorescent reporter could be used with the doxycycline inducible tRNA expression system components described above. The p413 TEF containing the dual fluorescent reporter was used as a template for Q5 mutagenesis PCR wherein the S151 mCherry residue, shown previously to be crucial to fluorescence (Rozik et al., 2022), was changed to various alternate codons corresponding to our mutant tRNA anticodon variants.

### Fluorescence microscopy and fluorescence intensity measurement

Yeast strains containing a combination of an inducible tRNA expression system plasmid and a mutant fluorescent rescue mistranslation reporter plasmid were grown overnight in the appropriate selective dextrose media at 30C and 250rpm orbital shaking. The next day, OD600 of overnight cultures was measured and a calculated volume of each culture was added to 3mL of selective dextrose media containing 1ug/mL of doxycycline such that an OD600 of 0.1 was achieved. Newly diluted cultures were incubated at 30C with 250rpm orbital shaking for 24 hours to allow ample opportunity for mistranslation of fluorescent proteins. A 10uL aliquot of ach culture was then transferred to a microscope for recording of fluorescence microscopy images. Photographs were taken using Nikon Eclipse Ci-L microscope, Nikon DS-Fi1camera and Nikon Elements software. Equivalent exposure time and gain were used for each image captured from the same sample across different fluorescent wavelengths.

Fluorescence of these cultures was also measured more directly by plate reader. Each culture was measured in quadruplicate for OD600, GFP signal, and RFP signal using a Tecan Infinite M200 plate reader and iControl software. Relative RFP/GFP signal was calculated relative to OD600 and compared to reference control ratios (wildtype mCherry construct co-expressed with wildtype tSerUGA1-1).

## Results

Previous iterations of this technique have used a GFP-mCherry fusion protein, with mCherry bearing a loss of function mutation, while the GFP acts as an expression control with which to compare RFP (plasmid WT-Pan, Addgene #99638)(Gomes et al., 2016; Sagi et al., 2016). Expression of this transgenics from the yeast expression plasmid used here produced no fluorescent signal from either protein, likely due to poor codon optimization of GFP in yeast cells, and thus poor translation. As a result, we adopt a yeast optimized eGFP (yoeGFP) linked to mCherry with a self-digesting T2A peptide (Addgene #118446)(Figure 1). Expression of this transgenic produces reliable GFP and mCherry signal in yeast cells (Figure 2, 3a).

**Figure 2.**
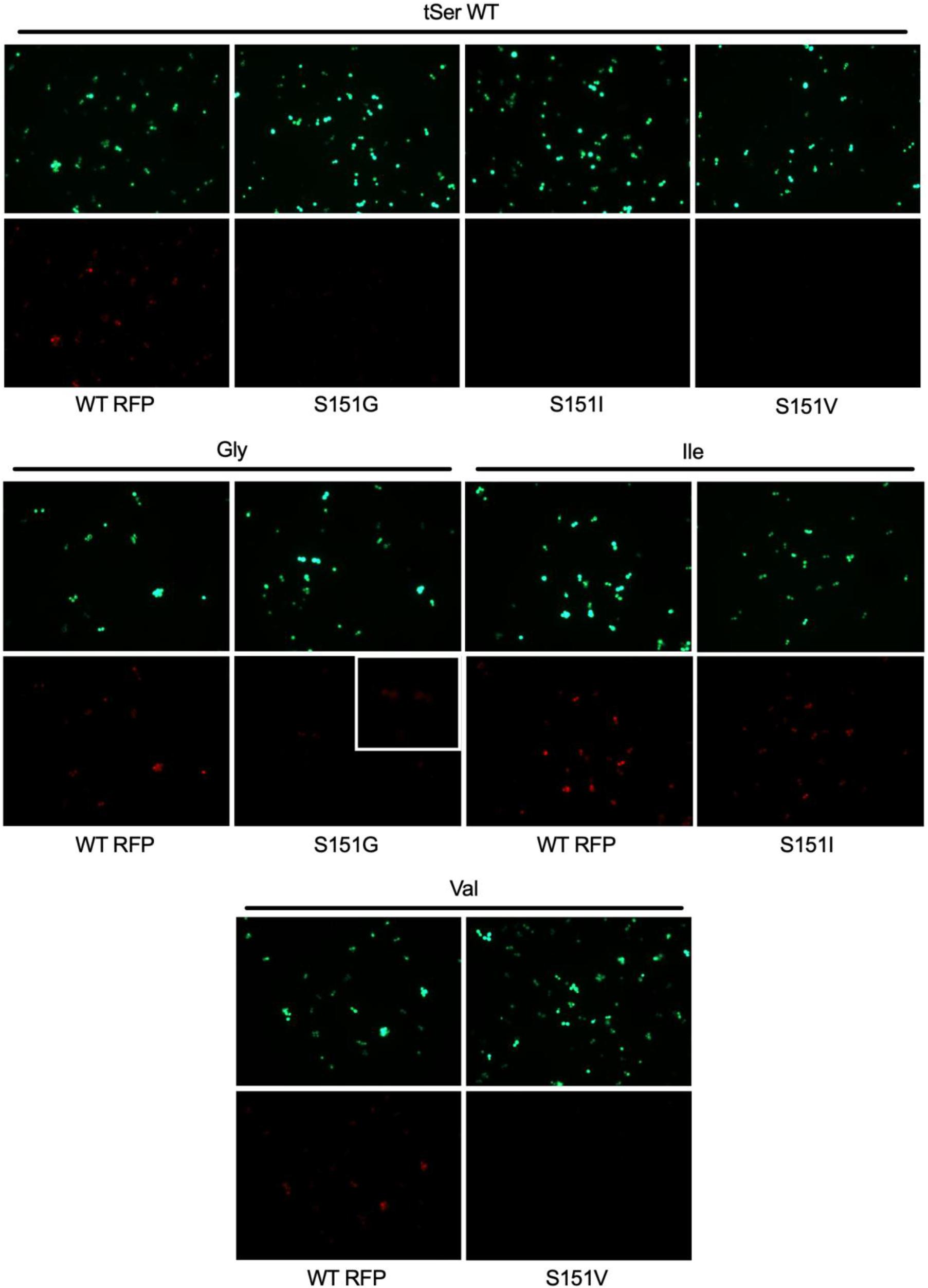
Fluorescence microscopy to detect mistranslation by rescue of mutant fluorescent proteins in yeast cells. GFP signal serves as an expression control with which to measure relative RFP signal. Top: expression of mCherry RFP mutants ablates relative RFP signal when expressed alongside wildtype tRNAs from the *Drosophila* tSerUGA1-1 gene. Expression of wildtype mCherry RFP serves as an RFP detection control. Middle and bottom: expression of mCherry mutants alongside corresponding rescuing forms of mistranslation. Some forms of mistranslation visibly partially rescue fluorescence of mCherry mutants (Ile and Gly). Expression of wildtype mCherry alongside mistranslating tRNA variants again serves as an RFP detection control.

**Figure 3.**
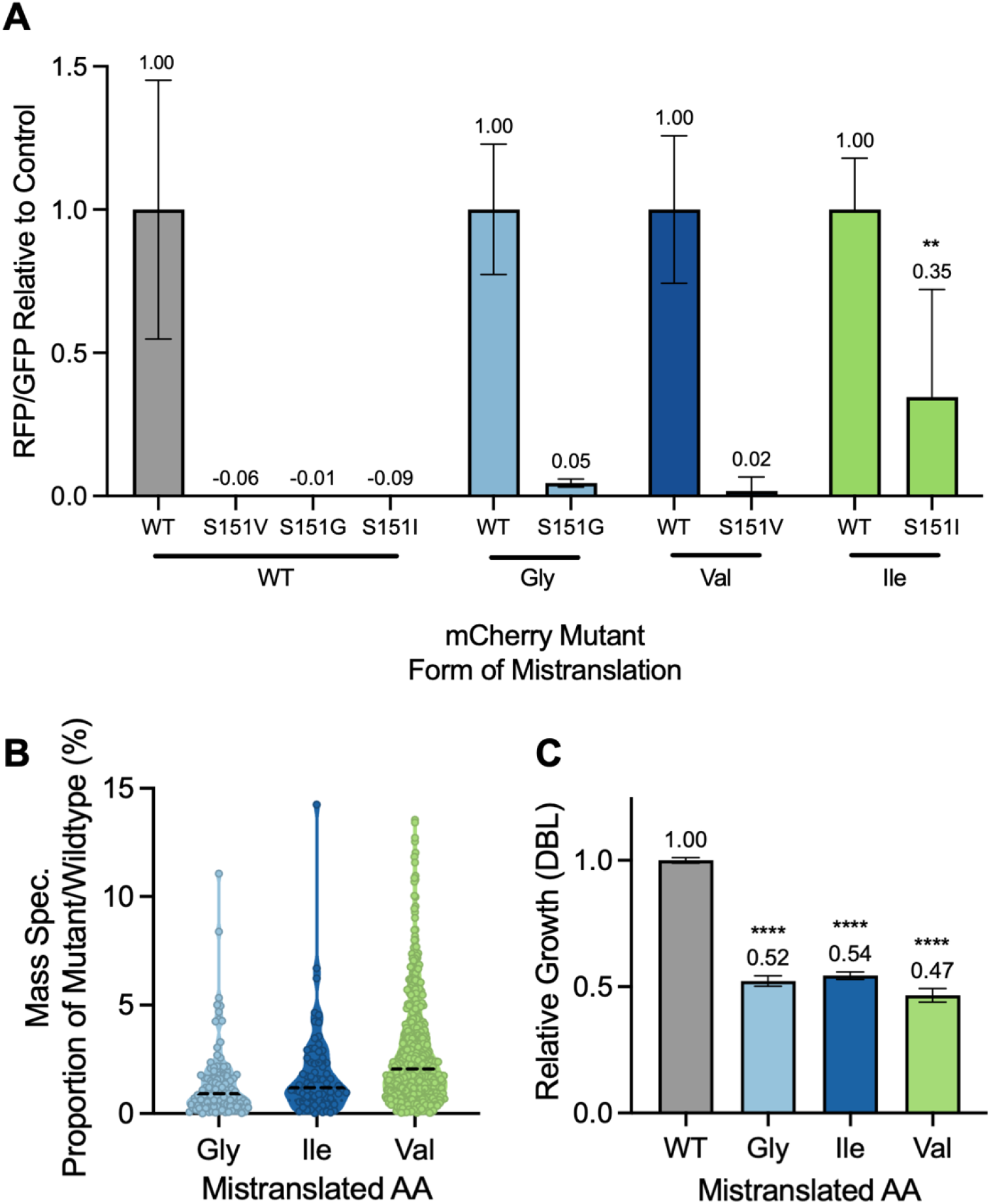
Mistranslation detection by plate reader measurement of fluorescence rescue. A) Y-axis measures RFP signal / GFP signal for each yeast strain relative to its appropriate tRNA control expressing a wildtpe mCherry RFP. GFP expression is used for relative RFP calculation to account for mistranslation / stress-induced reduction in transcription or translation. Each bar represents a unique combination of mCherry variant and tRNA variant. Groups expressing the same mistranslating tRNA variant are clustered along the X-axis and are filled with identical colours. Fluorescence of mCherry mutants is differentially restored by varied forms of mistranslation. Group means are listed as annotations above each bar (n=4). Statistically significant differences in quantity of RFP rescue by mistranslation relative to the appropriate WT tRNA^Ser^ control was determined by two-way ANOVA interaction effects (* denotes p<0.05). Error bars denote standard deviation. B) Mass spectrometry measurements of mistranslation quantity. Each dot represents the proportions of mistranslated / wildtype peptide as a percentage, with the mean proportion donated by black dashed line. C) Growth impairments imbued by mistranslating tRNA^Ser^ variants assessed. Growth is reported as doubling time relative to wildtype tRNA^Ser^ control. Data in B and C panels originally published in Charles & Moehring, 2026.

Mutagenizing the S151 residue of mCherry to various codons successfully ablates fluorescence (Figures 2,3a). Other residues crucial mCherry or yoeGFP function can also be mutagenized to assess mistranslation of different amino acids, such as mCherry A49 for alanine mistranslation (Rozik et al., 2022). Co-expression of these loss-of-function mutants with cognate mistranslating tRNA mutants restores mCherry signal to levels higher than those observed when no mistranslation is present (Figure 2, 3a). Plate reader quantification of GFP and mCherry signals confirms partial rescue of mCherry signal by some mistranslating mutants (Figure 3a). As expected, the RFP signal strength from successful rescue of mCherry differs with between mistranslated codons, likely owing to differences in tRNA pool competition. Interestingly, the strength of RFP rescue by each form of mistranslation differs substantially from predictions based on our previous measurements of mistranslation frequency by mass spectrometry (Figure 3B). Both methods for quantifying mistranslation appear disconnected from toxicity, as each form of mistranslation assessed are similar in the growth deficits imbued to yeast cells (Figure 1C).

## Discussion

These data demonstrate mistranslation fluorescence rescue as a cost-effective alternative to mass spectrometry in detecting mistranslation by mutant tRNAs in yeast. It is important to note that although only one form of mistranslation rescued RFP signal measured by plate reader to statistically significant levels (Ile), other forms of mistranslation are visible via fluorescence microscopy (Gly) and are measured at RFP signals above zero by plate reader. As such, mere detection of fluorescence rescue may be sufficient to confirm the occurrence of mistranslation. However, fluorescence rescue may have limited utility in providing accurate quantification of mistranslation frequency. While mass spectrometry reports Val-AAC mistranslation as producing the most mutant peptides of the strains assessed, the same Val-AAC mistranslation produces the weakest fluorescence rescue of the S151V mCherry variant. Given the well-established use of mass spectrometry in the field of mistranslation and the relative novelty of the specific iteration of fluorescence rescue via mistranslation, it would be fair to assume mistranslation frequencies identified by mass spectrometry are more reliable. If true, the yeast-specific fluorescence rescue reporter presented here is useful for the detection of the presence of mistranslation but may be inadequate for quantification of mistranslation.

However, it’s also possible that the differences in mistranslated product reported by each of these methods represents real biological differences in the proteins assessed. It may be the case that the 24 hours of cell growth allotted for mistranslated peptides to be produced is also sufficient time for several such mutant proteins to be degraded by various proteostatic cell stress responses, resulting in an overall lower abundance. For instance, given the high mRNA prevalence of Ile-AAT and the growth deficits observed for this form of mistranslation (Figure 3c), the mutant proteins produced may induce significant proteotoxic stress, therefore inciting significant action of stress response pathways that lead to the degradation of such proteins.

Through this lens, the number and frequency of mutant peptides identified via mass spec may represent only a portion of mistranslated proteins produced, and a second portion is detected and degraded by proteostatic mechanisms prior to measurement by mass spectrometry. The relative proportion mistranslated protein products that undergo pre-mass spectrometry clearance would vary with the ability for the cell to recognize and degrade the products of different forms of mistranslation.

Importantly, the degradation of mistranslated proteins prior to their detection is a concern unlikely to be shared by mistranslation fluorescence rescue. While many products of mistranslation may be degraded due to their misfolding, the mistranslation of mCherry at S151 does not cause such misfolding, but rather the restoration of conformation that allows its rescued fluorescence. Therefore, the fluorescence produced by mistranslation fluorescence rescue may hypothetically more accurately represent the total quantity of mistranslated protein, rather that the portion of mistranslated protein that evades degradation. This would be consistent with Ile-AAT mistranslation resulting in the highest fluorescent rescue signal of the forms of mistranslation assessed.

The ability to detect the presence of mistranslation can have particularly useful applications in examining tissue- or temporally-specific expression within *D. melanogaster* or other multicellular models of mistranslation. As such, the translation of this, similar, or alternative *Drosophila* codon-optimized dual fluorescent rescue reporters into flies warrants strong consideration for future works.

## Conclusion

The methods presented here allow for inexpensive detection of mistranslation in yeast cells. Given the frequent use of yeast as a model to research the effects of mutant tRNA mistranslation, the establishment of a yeast-specific iteration of this genetic technology presents high utility for future works. However, additional research is required to determine the accuracy of this approach’s quantification of mistranslation frequency and the source of its discrepancy between more commonly used mass spectrometry methods.

## Data Availability Statement

Plasmids are available upon request. The authors affirm that all data necessary for confirming the conclusions of the article are present within the article, figures, and supplemental material.

## Acknowledgements

We would like to thank Dr. Patrick O’Donoghue and Dr. Peter Rozik for their insight and donation of WT-Pan plasmids. We would also like to thank Dr. Kyle Hoffman and Bioinformatics Solutions for their efforts in collecting and analyzing the mass spectrometry experiments.

